# A pathogenic role for IL-10 signalling in capillary stalling and cognitive impairment in type 1 diabetes

**DOI:** 10.1101/2024.04.01.587630

**Authors:** Sorabh Sharma, Manjinder Cheema, Kelly A. Tennant, Roobina Boghozian, Ana Paula Cota, Tara P. Brosschot, Rachael D. Fitzpatrick, Jakob Körbelin, Lisa A. Reynolds, Craig E. Brown

## Abstract

Vascular pathology is associated with cognitive impairment in diseases such as type 1 diabetes, but precisely how capillary flow is affected and the underlying mechanisms remain elusive. Here we show that capillaries in the diabetic mouse brain are prone to stalling, with blocks composed primarily of erythrocyte plugs in branches off penetrating venules. Increased capillary obstructions were evident in both sexes and only partially reversed by insulin. Screening for circulating inflammatory cytokines revealed persistently high levels of interleukin-10 (IL-10) in diabetic mice. Contrary to expectation, stimulating IL-10 signalling increased capillary obstructions, whereas inhibiting IL-10 receptors with neutralizing antibodies or endothelial specific knockdown in diabetic mice, reversed these impairments. Chronic IL-10R blocking antibody treatment in diabetic mice also improved stimulus evoked cerebral blood flow, increased capillary widths in lower-order branches and reversed cognitive deficits. These data suggest IL-10 signalling plays an unexpected pathogenic role in cerebral microcirculatory defects and cognitive impairment.

## Introduction

The brain is the most metabolically demanding organ in the body, but lacks significant stores of energy ^1^. Consequently, the brain is heavily dependent on an uninterrupted supply of oxygen and nutrients from densely interconnected cerebrovascular networks that finely regulate blood flow to meet changes in metabolic demand ^2–4^. Capillaries in particular, play a crucial role in nourishing the brain as they represent >95% of the vessel length density ^5–7^ and their narrow, low pressure features maximize surface area and contact time of blood constituents for oxygen and nutrient exchange ^8^. The double edged sword of these features, however, is they also make capillaries especially prone to short and long-lasting stalling events, ranging from several seconds to days. *In vivo* imaging studies from our lab and others have revealed that at any given moment, 0.1-2% of capillaries in the healthy adult mouse brain are stalled ^9–11^. It is therefore not surprising that the incidence and duration of these stalling events dramatically increases with disease states such as ischemic or hemorrhagic stroke, Alzheimer’s disease, cytokine storms associated with cancer therapy or lipopolysaccharide treatment ^12–16^. Importantly, therapies targeting one of the root causes of these stalls, namely adherent neutrophils, leads to improved recovery after stroke or restores cognitive function in mouse models of dementia ^9, 14, 17, 18^.

Diabetes is one of the most prevalent chronic diseases in the world (∼422 million people affected) that is well known to produce vascular complications in the eye, heart, and kidney. In the brain, these complications manifest with increased incidence of stroke and substantially greater risk for cognitive impairment and dementia ^19–21^. Given these neurological complications, it is rather surprising that there has not been more extensive work characterizing the etiology and nature of microcirculatory perturbations in the diabetic brain. Indeed, most work thus far has focused on diabetes related changes in regional cerebral blood flow, myogenic tone of large cerebral arteries, blood brain barrier (BBB) permeability or susceptibility to damage after embolic occlusion of large vessels ^22–26^. Therefore, we sought out to understand whether capillary networks in a mouse model of type 1 diabetes (T1DM) were prone to stalling, what cells comprise these stalls and what molecular mechanisms could underlie these defects. In support of our hypotheses, we discovered that mice with T1DM had significantly higher rates of stalled capillaries in both sexes that were mostly occluded by red blood cells rather than leukocytes. Looking for mechanistic clues, we screened for multiple cytokines and chemokines in the blood and found abnormally high levels of interleukin-10 (IL-10) in diabetic mice. Contrary to initial expectations that IL-10 would serve a protective role as commonly reported in other conditions such as infection, stroke or aging ^27–31^, stimulating IL-10 signalling increased capillary obstructions/stalls in healthy mice whereas blocking IL-10 in diabetic mice helped prevent these microcirculatory disturbances and cognitive impairment. Collectively these findings strengthen the theory that microvascular pathology plays a significant role in cognitive decline and highlight the complex and often duplicitous roles that cytokines can play in neuro-inflammation depending on the biological context (infection, vs injury vs chronic autoimmune disease).

## Results

### Microcirculation in the diabetic brain is prone to stalling

There is considerable evidence indicating that type 1 diabetes exacerbates capillary plugging in peripheral organs, such as the heart, kidney and eye ^32–35^. However, it is unclear whether a similar phenomenon occurs in the brain. To address this question, we performed longitudinal *in vivo* two-photon imaging of the vasculature in the somatosensory cortex (typical x-y-z stack: 489×489×200µm) of adult non-diabetic mice or those treated with streptozotocin (STZ) to model type 1 diabetes (blood glucose >15mM; **Fig. 1a,b**). Under light isoflurane anesthesia, the blood plasma was labeled with intravascular injection of fluorescently tagged 70kDa dextrans and imaged from 4-8 weeks after the onset of hyperglycemia was confirmed (**Fig. 1a,b**). Since our steps in the axial plane were small enough that multiple images of the same capillary could be discerned over time, flowing capillaries were identified by the presence of red blood cell (RBCs) streaks moving through the vessel lumen. Stalled capillaries were defined as those with cells stuck in the vessel (dark gaps denoted by arrows in **Fig. 1c**) or those where no streaks for at least 3 image frames (minimum stall duration of 6.5s; **Fig. 1c**), similar to previously published criteria ^14^. As expected, repeated imaging of the same cortical areas over several weeks time revealed relatively low rates of stalled vessels in healthy mice (**Fig. 1d**; ∼90-94 stalls/mm^3^). However, diabetes significantly increased the number of stalled capillaries at each imaging time point (**Fig. 1d**; ∼258-374 stalls/mm3; Main effect of Diabetes F_(1,18)_=18.96, p<0.001).

**Figure 1.**
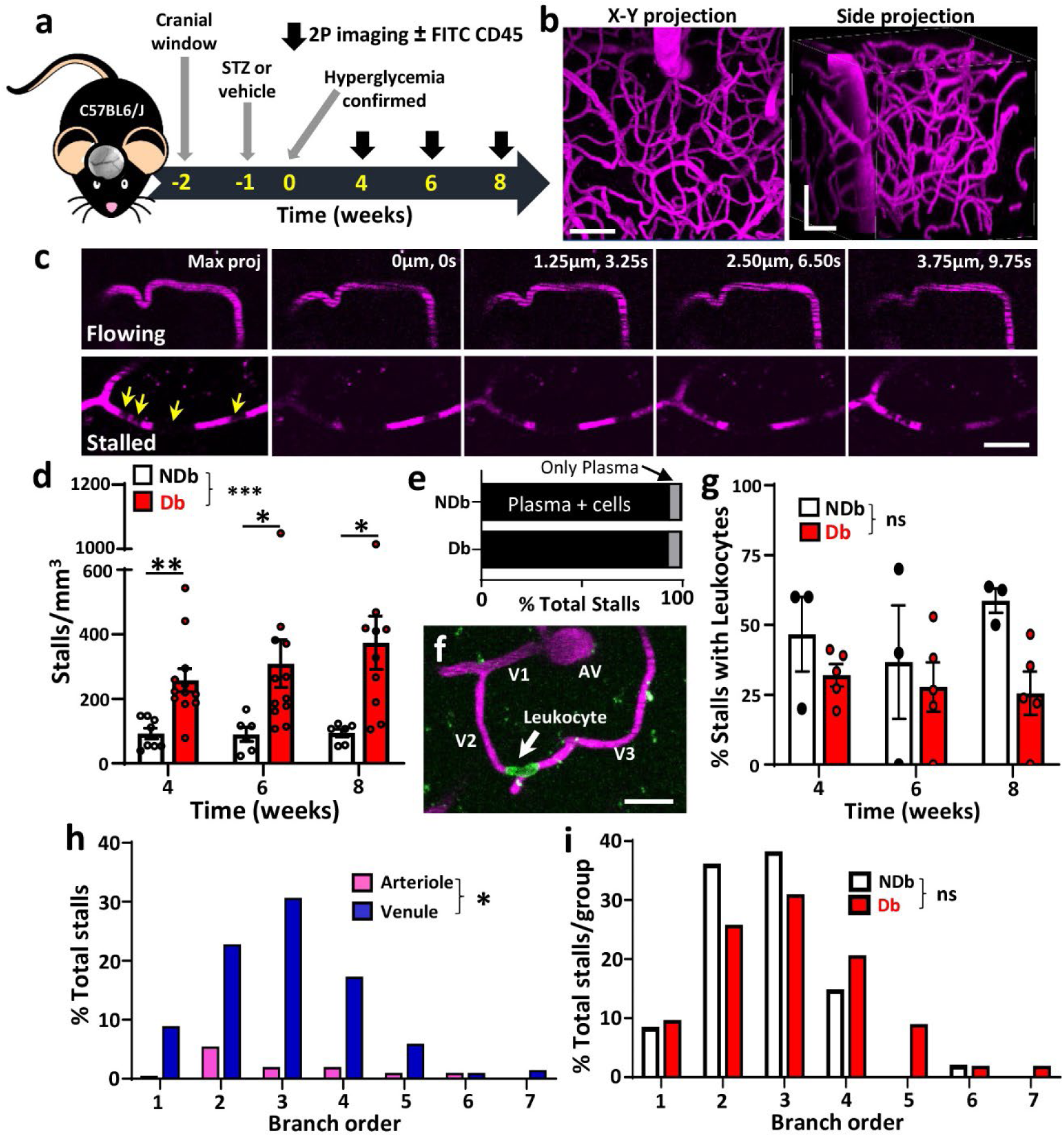
*In vivo* imaging shows abnormally high levels of stalled capillaries in diabetic mouse cortex. (**a**) Experimental timeline of diabetes induction and 2-photon imaging. (**b**) Maximal intensity two-photon X-Y and side (X-Z) projections showing cerebral vasculature (in magenta). (**c**) Successive images through a Z-stack show flowing or stalled capillaries. Note streaking pattern caused by RBC movement in flowing capillary or absence of streaking in those that are stalled. Yellow arrows indicate unlabelled cells stalled within the capillary. (**d**) Estimated stalls per mm^3^ in non-diabetic (n=6-8 mice) and diabetic mice (n=11-12 mice) in the weeks after hyperglycemia was confirmed. (**e**) Graph showing percentage of stalls with blood cells plugging the capillary or without (plasma + cells or just plasma). (**f**) Representative image showing a CD45.2 labeled leukocyte plugging a second order branch (V2) from a penetrating venule (PV). (**g**) Graph showing percentage of leukocyte stalled capillaries in non-diabetic and diabetic groups (n=3 and 5 mice, respectively) at different time points. (**h**) Percentage of stalled capillaries as it relates to branch order of the penetrating arteriole or venule. (**i**) Stalled capillaries as a function of arteriole or venous branch order in non-diabetic and diabetic groups. Abbreviations: NDb=Non-diabetic; Db= Diabetic; STZ= Streptozotocin. Data in d and g was analysed with Two-way ANOVA and Sidak multiple comparisons tests. Data in h-i was analysed with Mann-Whitney test. ns: not significant, *p<0.05, **p<0.01, ***p<0.001. Error bars represent mean ± SEM. Scale bars = 50µm (a) or 20µm (c,f).

To ascertain the characteristics of cerebral capillary stalls, we first classified stalls based on their appearance, i.e., whether they exhibited visible cells or not. The overwhelming majority of stalls in both experimental groups (93-94%) were those with cells plugging the capillary, whereas a small fraction did not appear to have cells in the lumen nor any evidence of streaking (termed “only plasma” in **Fig. 1e**). To determine the predominant cell type responsible for plugging capillaries (i.e., RBC or leukocytes), a subset of mice were imaged as described in **Fig. 1a**, and injected with FITC conjugated anti-CD45.2 antibodies to label circulating leukocytes^36^. As shown in **Fig. 1f**, leukocytes were easily visualized within stalled capillaries. While previous studies have identified leukocytes as the primary cause of stalls in a mouse model of Alzheimer’s disease or stroke ^14, 18^, our results indicate that only 25-32% of total stalls in diabetic mice were associated with a plugged leukocytes (**Fig. 1g**). Our statistical analysis revealed that the fraction of leukocyte stalled capillaries did not differ significantly between normal and diabetic groups, nor did it change over time (**Fig. 1g**; Main effect of Diabetes: F_(1,6)_=3.90, p=0.10; Main effect of Time: F_(1.06,6.37)_=3.90, p=0.46). Lastly, in order to understand what parts of the vascular tree were most susceptible, we examined where stalls occurred as a function of branch order from the penetrating arteriole or ascending venule (see branch orders from ascending venule in **Fig. 1f**). Our analysis shows that most stalls occur in the lower order capillary branches from the ascending venule (V2-4, **Fig. 1h**), which did not differ between diabetic and non-diabetic mice (**Fig. 1i**). Collectively, these results show that diabetic mice are prone to stalled capillaries that are mostly blocked by presumptive red blood cells (ie. cells not labelled with CD45), especially in capillary branches proximal to the ascending venule.

### Effect of insulin treatment and sex on capillary obstructions

To extend our *in vivo* observations and improve our understanding of how sex and insulin treatment affect capillary stalling in various brain regions, we intravenously injected 5µm diameter fluorescent microspheres in male and female diabetic mice treated with or without insulin and extracted their brain 3 days after injection (**Fig. 2a**). Using the microsphere assay allowed us to titer the density of microsphere obstructed capillaries to a level that was comparable to our *in vivo* estimates of stalled capillaries (∼30-150 microspheres/mm^3^ vs. 90-300 stalls/mm^3^). Previous work from our lab and others has shown that these small microspheres do not cause blood brain barrier disruption, local recruitment of microglia or micro-infarcts that are commonly found after injection of larger emboli (>10µm diameter) ^5, 10, 37, 38^. Furthermore, since the dose of microspheres can be carefully controlled between mice, we could perform a high throughput screen of obstructed capillary density across many experimental groups and different brain regions not easily accessible with *in vivo* microscopy ^5, 10^. As anticipated, STZ treatment led to a significant increase in blood glucose levels in the uncontrolled diabetic group (F_(1,114)_=176.1, p<0.0001), whereas insulin treatment lowered blood glucose but not to completely normal levels (F_(1,65)_=32.8, p<0.0001; compare “Db” and “Db + Ins” to “NDb” groups in **Fig. 2b**). Relative to non-diabetic control mice, body weights were lower for both uncontrolled (F_(1,114)_=17.55, p<0.0001) and insulin treated diabetic mice (F_(1,65)_=10.21, p<0.01) although this was not statistically significant at any specific time point (**Fig. 2c**). Consistent with our *in vivo* findings on capillary stalling, we found a significant increase in the number of microsphere obstructed capillaries in the forebrain of diabetic mice, when collapsed across sex (upper panel in **Fig. 2d**). Insulin treatment did not significantly lower the density of obstructions from that found in uncontrolled diabetic mice (**Fig. 2d**). Consistent with a recent study ^39^, plotting blood glucose values in all groups as a function of obstruction density revealed a significant positive linear relationship (**Fig. 2e**; R^2^=0.52, p<0.0001). Based on previous work showing there are significant regional differences in susceptibility to capillary stalling ^5^, we also quantified microspheres in different brain regions. This analysis indicated that the density of capillary obstructions was significantly greater in uncontrolled diabetic mice relative to non-diabetic controls (**Fig. 2f**; Effect of Diabetes: F_(1,373)_=59.1, p<0.0001), which also varied significantly by brain region (Effect of Region: F_(12,373)_=20.31, p<0.0001; Diabetes x Region interaction: F_(12,373)_=1.7, p=0.06) with highest density in retrosplenial cortex. Insulin treated diabetic mice showed a slight reduction in obstruction density compared to uncontrolled diabetic mice (**Fig. 2f**; Effect of Insulin: F_(1,244)_=4.39, p=0.04). Lastly, we stratified our obstruction density analysis by sex, and discovered there were no sex differences in the density of obstructed capillaries in non-diabetic or diabetic mice (**Fig. 2g**; Effect of sex in NDb mice: F_(1,181)_=0.66, p=0.42 and Db mice: F_(1,165)_=0.25, p=0.62). These results indicate that increased levels of capillary obstructions in diabetic mice are only partially affected by insulin treatment and not dependent on sex.

**Figure 2.**
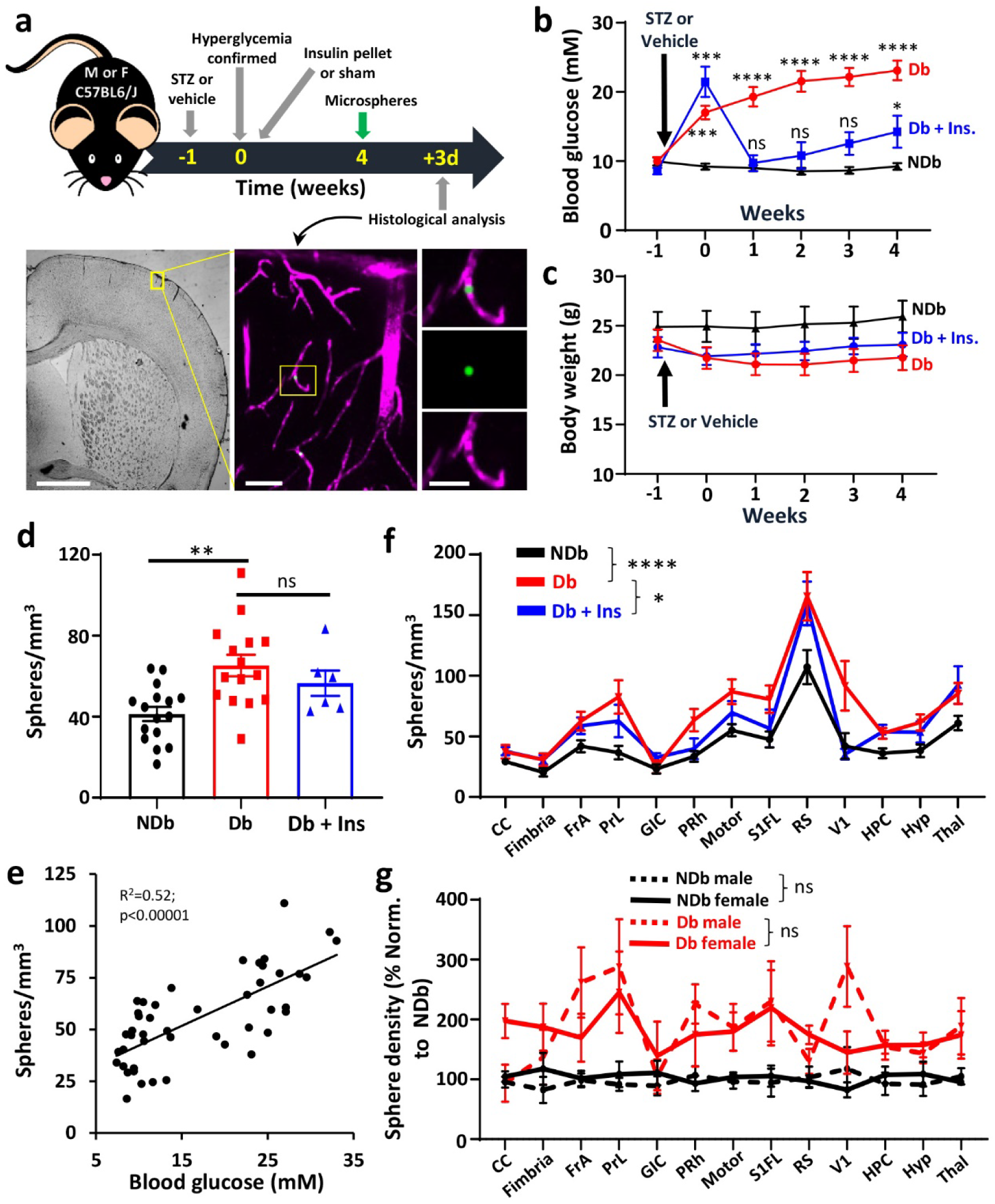
Diabetic mice exhibit more plugged capillaries which is not affected by sex and only partially ameliorated with insulin treatment. (**a**) Top: Timeline of diabetes induction and microsphere injection experiments. Bottom: Widefield images showing fluorescent microspheres (green) plugging the lumen of capillaries filled with fluorescent dextrans (magenta) in the somatosensory cortex. (**b**) Weekly blood glucose levels in experimental groups. (**c**) Weekly body weight measurements in experimental groups. (**d**) Quantification of microsphere plugged capillaries across forebrain in non-diabetic, uncontrolled diabetic and insulin treated mice (n=16, 15 and 6 mice, respectively). (**e**) Regression analysis showing significant relationship between blood glucose levels and forebrain density of microsphere plugged capillaries (n=37 mice). (**f**) Density of microsphere plugged capillaries across different brain regions in each experimental group. (**g**) Regional density of microsphere plugged capillaries as a function of sex (NDb: n=8 male and female mice; Db: 7 male and 8 female mice). Abbreviations: **NDb:** Non-diabetic, **Db:** Diabetic, **Db+Ins:** Diabetic + insulin treatment. **CC:** Corpus callosum; **FrA:** Frontal association cortex; **PrL:** Prelimbic cortex; **GIC:** granular/dysgranular insular cortex; **PRh**: Peri-rhinal cortex; **Motor:** primary/secondary motor cortex; **S1FL:** primary forelimb somatosensory cortex; **RS**: retrosplenial cortex; **V1:** primary visual cortex; **HPC:** hippocampus; **Hyp:** Hypothalamus; **Thal**: Thalamus. Data in b, c, f and g analysed with two-way ANOVA followed by Tukey’s multiple comparison test when appropriate. Data in d analysed with two-tailed unpaired t-test. ns: not significant, *p< 0.05, **p<0.01, ***p< 0.001, ****p<0.0001. Error bars: mean ± SEM. Scale bars in a, from left to right = 1mm, 50µm and 20µm.

### Persistently high levels of circulating IL-10 in diabetic mice is associated with obstructed capillaries

To elucidate the mechanisms underlying the increased incidence of capillary stalling in diabetic mice, we screened for multiple cytokines/chemokines in blood serum samples collected after 8 weeks of uncontrolled diabetes using a multiplex assay. Our analysis revealed that levels of certain cytokines, including IL-1β, IL-2, IL-6, IL-10, IP-10, TNFα, KC-CXCL1, MIP-1,-2,-3 and MCP-1, were significantly higher in diabetic mice than in non diabetic control mice (**Fig. 3a**, for raw values see **Supp. Fig. 1a**). Given that diabetic mice show significantly higher rates of capillary stalling well before 8 weeks time (see **Fig. 1d**), we re-examined select cytokines IL-10, MCP1/CCL2, KC/CXCL1, IL-17 and MIG1/CXCL9 at 4 weeks. Among the cytokines re-examined, only IL-10 was significantly elevated at both 4 and 8 week time points relative to non-diabetic mice (**Fig. 3b** and **Supp. Fig. 1b-e**). This finding was surprising given that IL-10 is generally considered an anti-inflammatory cytokine; however recent reports shown that it can also have pleiotropic effects and function both as a pro-inflammatory and anti-inflammatory cytokine ^40^. To help us understand what effect elevated IL-10 could produce, we injected non-diabetic mice with IL-10 or control albumin protein (0.5µg, i.v.) and quantified the density of microsphere obstructed capillaries (**Fig. 3c**). Our results show that injecting IL-10 into the bloodstream can significantly increase capillary obstructions (**Fig. 3d**). These results suggest that circulating IL-10 could play a role in capillary stalling.

**Figure 3.**
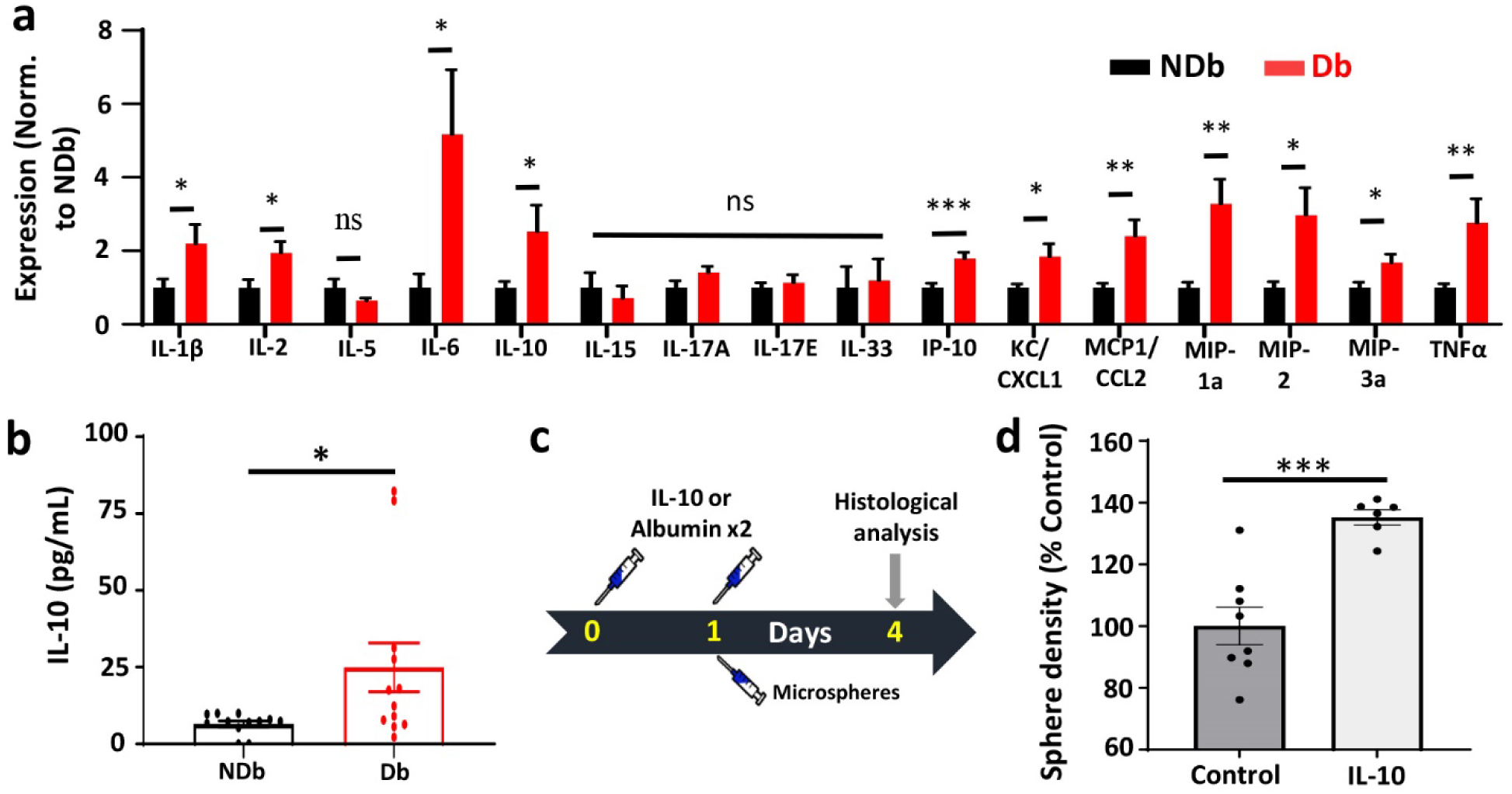
Elevated levels of IL-10 in diabetic blood is associated with capillary plugging. (**a**) Multiplex immunoassay shows normalized expression of cytokines and chemokines in blood serum of non-diabetic (n=12 mice) and diabetic mice (n=10 mice, collected 8 weeks after confirmation of hyperglycemia). (**b**) IL-10 levels are elevated in diabetic blood serum 4 weeks after confirmation of hyperglycemia. (**c**) Experimental timeline for examining the effect of intravenous injection of IL-10 or control albumin protein on capillary plugging. (**d**) Normalized density of microsphere plugged capillaries across forebrain in control and IL-10 injected groups (n=8 and 6 mice, respectively). Abbreviations: **IL-1β; -2; -5;-6; -10; -15; - 17a; -17e, -33**: Interleukin-1β; -2; -5;**-**6; -10; -15; -17a; -17e, -33; **IP-10**: Interferon gamma-induced protein 10; **KC/CXCL1:** keratinocyte chemoattractant/ chemokine (C-X-C motif) ligand 1; **MCP-1/CCL2:** Monocyte chemoattractant protein-1/ Chemokine (CC-motif) ligand 2; **MIP-1a:** Macrophage inflammatory protein-1 alpha; **TNF-α:** Tumour Necrosis Factor-α). Data in a, b and d analysed using two-tailed unpaired t-tests. ns: not significant, *p<0.05, **p<0.01, ***p<0.001. Error bars: mean ± SEM.

### Endothelial, but not neutrophil specific knockdown of IL-10 receptor signalling alleviates capillary obstructions in diabetic mice

Given that high levels of circulating IL-10 were associated with stalled capillaries and IL-10 receptors are expressed on both endothelial cells (**Supp. Fig. 2b**) and neutrophils, we next explored whether broad spectrum inhibition of the IL-10 receptor with a neutralizing antibody, or cell specific knockdown of IL-10 receptors in vascular endothelial cells or neutrophils, could prevent capillary obstructions in diabetic mice (**Fig. 4a,b**). To knockdown IL-10 receptors in endothelial cells, diabetic IL10ra floxed mice (*IL10ra^fl/fl^*) were intravenously injected with AAV-BR1-iCRE that selectively targets brain endothelial cells ^41^ or control vector (AAV-BR1-eGFP). For neutrophil specific knockdown, we crossed *Mrp8cre* mice with *IL10ra^fl/fl^* mice. Cre recombinase activity in either brain endothelial cells or neutrophils was confirmed in the cre-dependent tdTomato reporter mouse (Ai9; **Fig. 4c,d**). Furthermore, we verified *IL10ra* knockdown after AAV-BR1-iCRE injection from isolated endothelial cells using qPCR or PCR for *Mrp8cre:IL10ra^fl/fl^* mice (**Supp. Fig. 2**).

**Figure 4.**
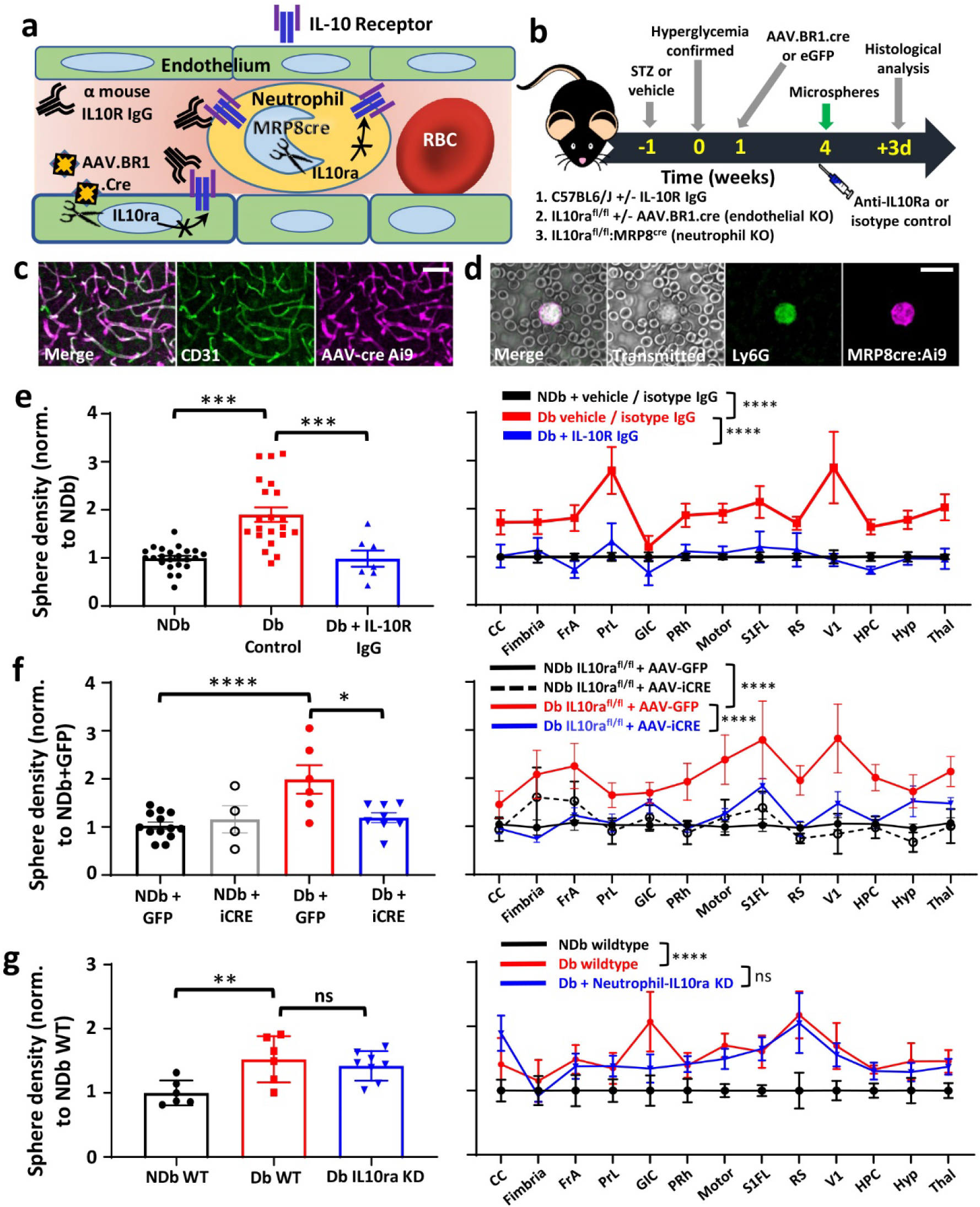
IL-10 receptor inhibition with neutralizing antibodies or endothelial specific knockdown alleviates capillary plugging in diabetic mice. (**a**) Illustration shows how IL-10 receptor signalling could impact capillary obstructions through vascular endothelial cells and/or neutrophils and therapeutic approaches tested. (**b**) Experimental timeline and 3 treatment groups for inhibiting IL-10 receptor signalling: i) injection of IL-10R neutralizing antibodies in C57BL6/J wild-type mice; ii) injection of AAV.BR1.iCRE or control virus (AAV.BR1.eGFP) in IL10ra^fl/fl^ mice for endothelial specific knockdown of IL10ra receptors, iii) neutrophil specific knockdown using IL10ra^fl/fl^ mice crossed with MRP8cre mice. Validation of Cre recombinase activity in either brain endothelial cells using CD31 immunolabelling (**c**) or neutrophils using Ly6G antibody (**d**) with cre-dependent tdTomato reporter mouse (Ai9). (**e**) Effect of IL-10R neutralizing antibody treatment on the density of microsphere plugged capillaries in non-diabetic controls, diabetic controls or diabetic + IL-10R IgG treated mice (n=23, 21 and 7 mice, respectively). Microsphere densities are shown collapsed across all forebrain regions (Left) or within specific brain regions (Right). (**f**) Effect of Endothelial specific IL10ra knockdown on density of plugged capillaries across the forebrain (Left) or within specific brain regions (Right) in non-diabetic (AAV-GFP: n=13 mice; AAV-iCre: n=4 mice) and diabetic mice (AAV-GFP: n=6 mice; AAV-iCre: n=8 mice). (**g**) Density of microsphere plugged capillaries in forebrain (Left) or specific brain regions (Right) in non-diabetic wildtype, diabetic wildtype or IL10ra knockdown in neutrophils (n=6, 6 and 8 mice, respectively). Data in e-f analysed with two-tailed unpaired t-tests (left panel) or two way ANOVA followed by Sidak’s multiple comparison tests (right panel). ns: not-significant, *p<0.05, **p<0.01, ***p<0.001, ****p<0.0001. Scale bars in c,d = 20µm. Error bars: mean ± SEM.

For broadly inhibiting IL-10 signalling, we treated hyperglycemic diabetic mice with intravenous injection of IL-10R neutralizing antibody or isotype control antibody. Mice were injected with microspheres and the density of obstructed capillaries was quantified 3 days later (see experimental timeline in **Fig. 4b**). Our results show that diabetic mice treated with the IL-10R neutralizing antibody had significantly lower density of obstructed capillaries in forebrain or across different brain regions relative to control treated diabetic mice (**Fig. 4e**; F_(1,335)_=32.83, p<0.0001). We next examined the effect of AAV mediated knockdown of endothelial IL-10 signalling. Diabetic *IL10ra^fl/fl^* mice injected with AAV-BR1-iCRE exhibited a significant reduction in the density of obstructed capillaries when compared to diabetic mice treated with control AAV-BR1-eGFP (**Fig. 4f**). This reduction was evident for forebrain analysis as well as across different brain regions (**Fig. 4f**; F_(1,156)_=48.10, p<0.0001). We next examined the density of obstructed capillaries in diabetic mice with neutrophil specific knockdown of IL-10 receptors or Cre negative (“wildtype”) controls (**Fig. 4g)**. Our analysis revealed no significant differences between diabetic treatment groups (**Fig. 4g**; F_(1,156)_=0.95, p=0.33). Importantly, none of the manipulations described above significantly altered blood glucose levels in diabetic mice, thereby arguing against the possibility that the rescue effects of IL-10R antibodies or *IL10ra* knockdown were caused by lowering blood glucose levels (**Supp. Fig. 3**; F_(5,38)_=0.53, p=0.75). In summary, these findings show that neutralizing antibodies or endothelial specific inhibition of IL-10 receptor signalling alleviate capillary obstructions in diabetic mice.

### Neutralizing IL-10 signalling alleviates capillary stalling *in vivo* and improves cerebral blood flow

Since treating diabetic mice with a neutralizing antibody represents the most clinically translatable approach for modulating IL-10 signalling *in vivo*, we next imaged the cerebral vasculature in diabetic mice after injection of isotype control antibody and then again 1 week later after treatment with IL-10R neutralizing antibody (**Fig. 5a**). Two-photon imaging of the cortical microcirculation indicated a significant decrease in the number of stalled capillaries, which amounted to a 63% reduction in stalling (**Fig. 5b**). In order to better understand how our treatment was affecting the microcirculation, we measured the width of capillaries across lower order branches (branch orders 1-4) from the penetrating arteriole or ascending venule (see examples in **Fig. 5a**). Plotting capillary widths indicated wider capillaries from lower order branches off the penetrating arteriole after IL-10R neutralizing antibody treatment (**Fig. 5c**). By collapsing width measurements across first to forth order capillaries on the arteriole or venule side, we found a significant increase in the diameter of capillaries branching off the penetrating arteriole after IL-10R neutralizing antibody treatment (**Fig. 5d**), whereas those on the venule side did not change significantly (**Fig. 5e**). These results indicate that IL-10R neutralizing antibody treatment reduces the fraction of stalled capillaries and dilates lower order capillaries off the penetrating arteriole.

**Figure 5.**
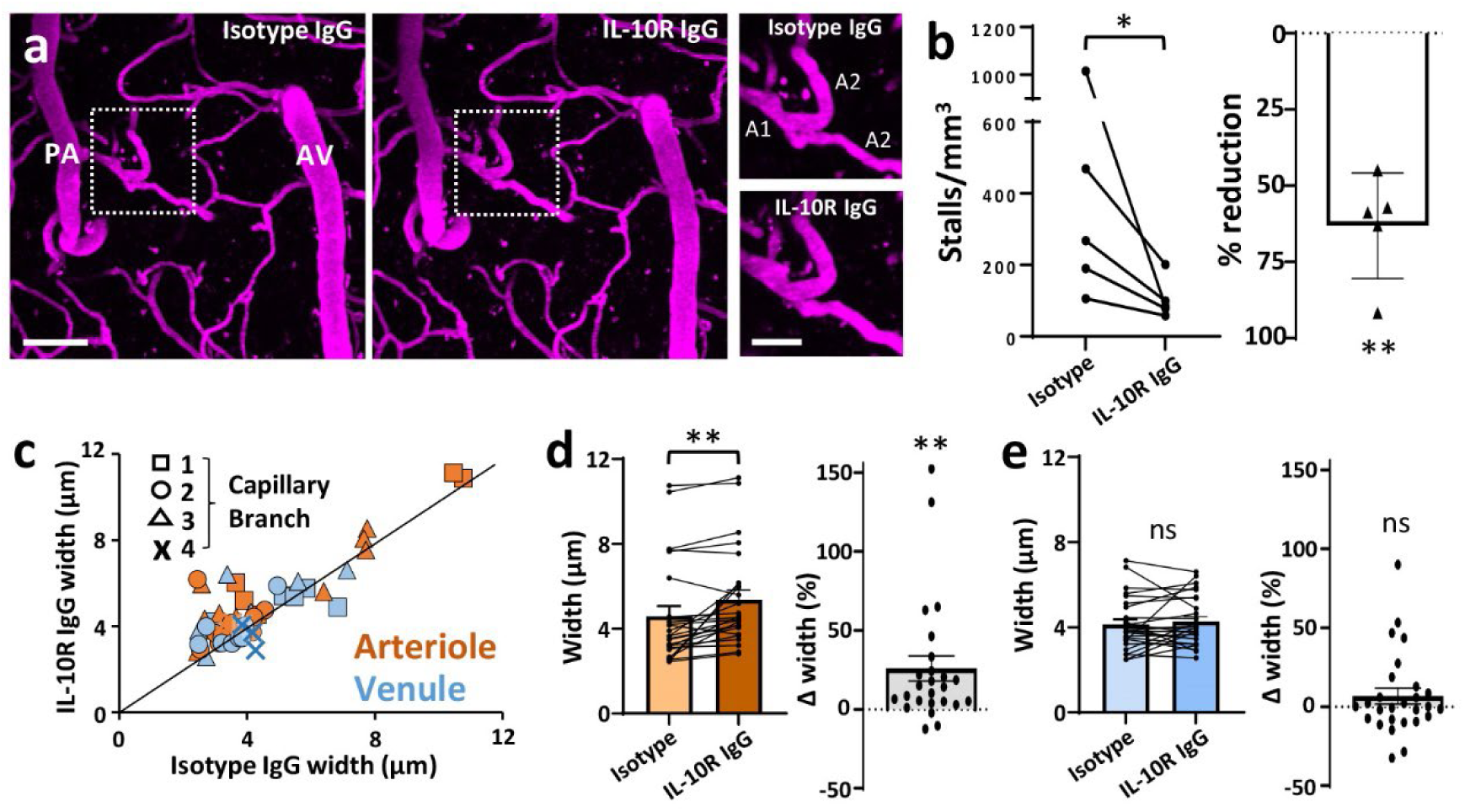
Treating diabetic mice with IL-10R neutralizing antibody lowers stalling rates and increases capillary width. (**a**) *In vivo* two photon image projections showing plasma labeled vasculature (in magenta) in the diabetic somatosensory cortex following injection of isotype control antibody and after treatment with IL10R neutralizing antibody one week later. Inset shows lower order capillary branches off the penetrating arteriole (PA) in each treatment condition. images of vasculature. (**b**) Graphs show absolute reduction in stalling density (left) or % change (right) in diabetic mice treated with isotype control or IL-10R neutralizing antibody (n=5 mice imaged after both treatments). (**c**) Unity plot shows width of lower order capillary branches (branches 1-4) off the penetrating arteriole (brown, n=25 capillaries) or ascending venule (blue, n=27 capillaries) following Isotype control or IL10R neutralizing antibody treatment. (**d**) Absolute or % change in width of capillaries branching off the penetrating arteriole. (**e**) Absolute or % change in width of capillaries branching off the ascending venule. Data in b, d, e analysed with two-tailed paired t-tests (left panels) or one-sample t-tests (right panels). PA: penetrating arteriole, AV: ascending venule, ns: not significant, *p<0.05, **p<0.01. Scale bars in a = 50µm, inset = 20µm. Error bars: mean ± SEM.

Next we examined whether chronic treatment of diabetic mice with IL-10R neutralizing antibodies could improve vascular responses to stimuli well known to increase blood flow (see **Fig. 6a**). Therefore we utilized laser doppler flowmetry to assess cerebral blood flow (CBF) in urethane anesthetized mice before and after 5% CO_2_ challenge, vibrotactile sensory stimulation of the limb or exposure to isoflurane. Our findings indicate that the normal increase in CBF following CO_2_ exposure was blunted in isotype treated diabetic mice, whereas treatment with IL-10R neutralizing antibody helped partially normalize CBF responses (**Fig. 6b-e**). Similarly, CBF responses to sensory limb stimulation were lower in isotype treated diabetic mice relative to non-diabetic mice (**Fig. 6f-i**). Treatment with IL-10R neutralizing antibody produced a small but significant increase in CBF responses, primarily near the end of the vascular response (7.5-15s after stimulation; **Fig. 6f-i**). Interestingly, no significant differences were observed among experimental groups in response to isoflurane stimulation (**Figure 6j-m**). This latter finding was important for ruling out the possibility that elevated rates of capillary stalling observed in diabetic mice *in vivo* were related to our use of isoflurane anesthesia during 2-photon imaging.

**Figure 6.**
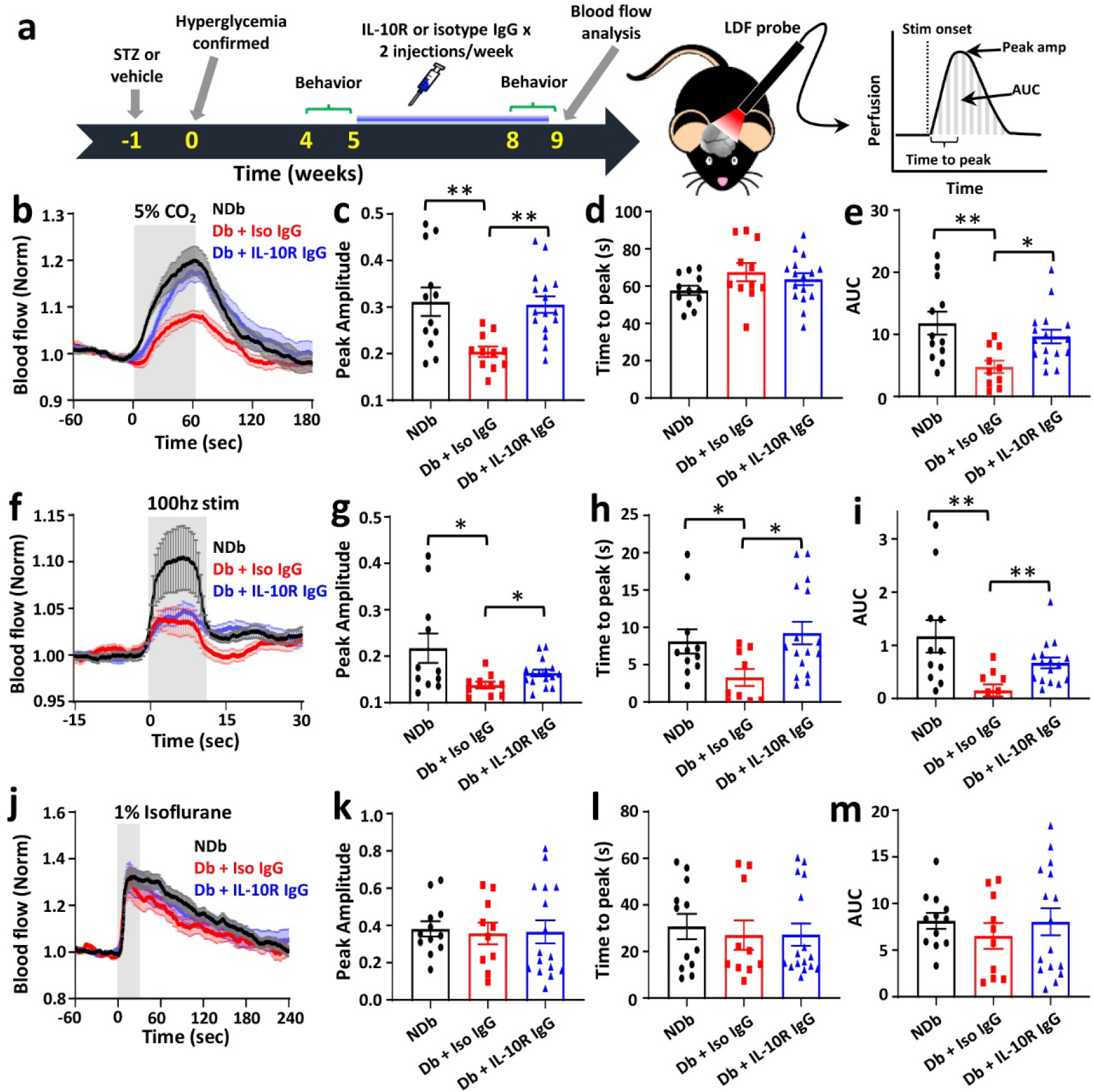
Effect of IL-10R neutralizing antibody on cerebral blood flow in diabetic mice. (**a**) Schematic showing timeline of experimental procedures and cerebral blood flow (CBF) measurements using laser doppler flowmetry (LDF). CBF measurements include peak amplitude, time to peak and area under the curve (AUC). (**b**) Blood flow changes in response to inhalation of 5% carbon dioxide (CO_2_). (**c-e**) Peak amplitude, Time to peak and AUC, respectively in non-diabetic or diabetic mice treated with isotype control or IL10R neutralizing antibody (n=12, 11 and 16 mice, respectively). (**f**) Blood flow changes in response to 100hz vibrotactile stimulation of the limb. (**g-i**) Peak amplitude, time to peak and AUC, respectively. (**j**) Blood flow changes in response to 1% Isofluorane mixed in air. (**k-m**) Peak amplitude, time to peak and AUC of blood flow changes, respectively. Grey areas in b, f and j indicate duration of blood flow stimuli (60, 10 and 30s, respectively). Data in c-e, g-i, k-m analysed by two-tailed unpaired t-tests. ns: not significant, *p<0.05, **p<0.01. Error bars: mean ± SEM.

### IL-10R neutralizing antibody treatment improves cognitive function

There is a growing body of evidence showing that vascular pathology plays a critical role in cognitive impairment ^42–44^. Since type 1 diabetes is associated with cognitive impairment, especially if poorly controlled^26, 45^, we tested whether chronic treatment with IL-10R neutralizing antibody could ameliorate deficits in cognitive and sensorimotor functions (see **Fig. 6a** for timeline). First we employed the novel object recognition (NOR), a common test of memory ^46^. This test showed that diabetes related impairments in the frequency of visits to novel zone (**Fig. 7a**) or time spent exploring the novel object (**Fig. 7b**), can be reversed in diabetic mice given IL-10R neutralizing antibody. We next assessed spatial learning and memory in the Morris water maze. At both 4 and 8 weeks time, control treated diabetic mice took significantly longer to find the submerged platform during the initial platform learning trials (**Fig. 7c**), as well as after the platform location was switched to a new location (“reversal learning” in **Fig. 7d**). In both tests, cognitive function improved in diabetic mice treated with IL-10R neutralizing antibody given they took significantly less time to find the platform relative to control treated diabetic mice (**Fig. 7c,d**). Performance in the memory probe trial was marginally worse in diabetic mice, and IL-10R neutralizing antibody treatment slightly, but not significantly increased this metric (**Fig. 7e**). It should be noted that group differences in learning and memory cannot be explained by differences in swim speed or gross visual abilities (based on the visible platform test) since no significant differences were found across experimental groups (**Supp. Fig. 4**). To determine if sensory-motor behaviours were affected, we examined performance in the adhesive tape removal test and open field activity. Control treated diabetic mice exhibited significantly longer latencies to remove the tape relative to IL-10R blocking antibody treated mice (**Fig. 7f**). While diabetic mice showed reduced activity in the open field, particularly at the 8 week time point, inhibiting IL-10R had no effect on this behaviour (**Fig. 7g**). Similarly in the elevated plus maze, anxiety-like behaviour in diabetic mice reflected by the lower preference for entry into open arms, was not affected by IL-10R treatment (**Fig. 7h**). In summary, these results show that IL-10R receptor inhibition in diabetic mice generally improves metrics of cognitive and somatosensory-motor function (NOR, water maze and tape test), but has minimal or no effect on general ambulatory or anxiety-like behaviours.

**Figure 7.**
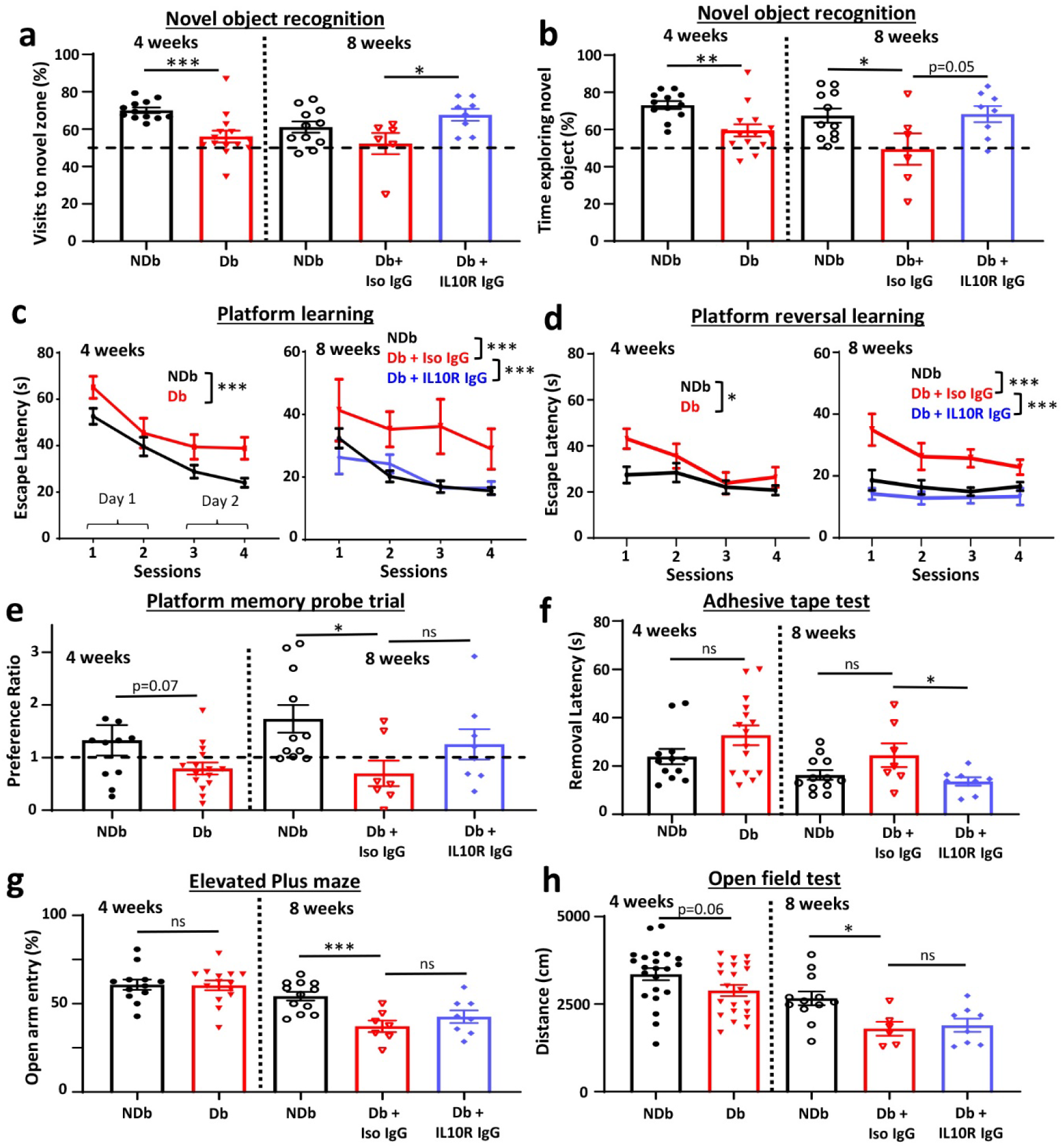
Long-term IL-10 receptor inhibition improves cognitive function in diabetic mice. (**a,b**) Graphs show the frequency of visits to novel object zone (a) or % time exploring novel object (b) at 4 weeks in non-diabetic or diabetic mice (n=12 and 14 mice, respectively), or 8 weeks in non-diabetic or diabetic mice treated with isotype control or IL10R neutralizing antibody (n=12, 6 and 8 mice, respectively). (**c**) Escape latency for learning hidden platform location in Morris water maze at 4 weeks in non-diabetic and diabetic mice (n=26 and 23 mice, respectively) or 8 weeks (n=12, 7 and 8 mice for non-diabetic, diabetic + Isotype and diabetic + IL10R neutralizing antibody, respectively). (**d**) Escape latency for learning new platform location in water maze (“platform reversal learning”) in non-diabetic and diabetic mice at 4 weeks (n=12 and 15 mice, respectively) and 8 weeks time (n=12, 7 and 8 mice for non-diabetic, diabetic + Isotype and diabetic + IL10R neutralizing antibody, respectively). (**e**) Preference for quadrant with hidden platform (“platform memory probe trial”) in non-diabetic and diabetic mice at 4 weeks (n=12 and 15 mice, respectively) and 8 weeks (n=12, 7 and 8 mice for non-diabetic, diabetic + Isotype and diabetic + IL10R neutralizing antibody, respectively). (**f**) Graph showing latency of tape removal (in seconds) in different experimental groups (n=12 and 15 mice at 4 weeks; n=12, 7 and 8 mice at 8 weeks). (**g**) % time spent in open arm on elevated plus maze for experimental groups (n=12 and 15 mice at 4 weeks; n=12, 7 and 8 mice at 8 weeks). (**h**) Distance travelled by mice in each experimental group in open field test (n=22 and 21 mice at 4 weeks; n=12, 6 and 8 mice at 8 weeks). Two-tailed unpaired t-tests were used to analyse data at each time point in a, b, e-h. Data in c, d were analysed with two-way ANOVA followed by Tukey’s multiple comparison tests. *p<0.05, **p<0.01, ***p<0.001. Error bars: mean ± SEM.

## Discussion

T1DM has been reported to increase the risk of neurological complications and lowers functioning in several cognitive domains ^20, 26, 47^. These diverse problems likely reflect the fact that diabetes disrupts micro-vascular networks throughout the body, including the brain ^48^. Therefore the goals of the present study were to examine the nature of microcirculatory disruptions in the diabetic brain, what molecular mechanisms and cell types contributed to this pathology and finally, what treatments could mollify the deleterious effects of diabetes on cognitive function. Here we demonstrate using longitudinal *in vivo* two-photon microscopy that T1DM in both male and female mice is associated with a 2-4 fold increase in stalled or obstructed cortical capillaries relative to healthy controls (eg. 8 wks: 364 vs 94 stalls/mm^3^). Of note, treating diabetic mice with insulin was not sufficient to fully normalize the risk of capillary obstructions, which is consistent with other studies showing that partial normalization of blood glucose levels does not necessarily rescue diabetes related neuropathology ^49–52^. Assuming there are approximately 10,000-20,000 capillaries per mm^3^ ^10, 53^ our experiments would suggest about 0.465-0.93% of capillaries in healthy mice or 1.82-3.64% in diabetic mice are stalled at any given time. These estimates closely match other *in vivo* studies in healthy mice, including those using awake imaging preparations, and slightly higher rates in diabetic mice relative to that found in mouse models of Alzheimer disease ^9,12, 14^. Furthermore, the majority of stalls were evident in the 1-4^th^ order branches off penetrating venules, which agrees with previous work ^9, 14^.

In order to elucidate what type of blood cells cause capillary stalls, we fluorescently tagged leukocytes using a CD45.2 antibody. Previous work has shown that this antibody labels 95% of circulating leukocytes and does not cause any cell depletion, even at much higher doses than that used in the present study ^36^. Our experiments revealed that most stalls in diabetic mice were made of erythrocytes rather than leukocytes, thereby arguing that adherent leukocyte were not the primary driver of stalled capillaries. This finding contrasts with studies in the diabetic retina, where monocytes and neutrophils were abundantly found in capillaries ^54, 55^. However, it should be noted that these post-mortem studies were also associated with significant tissue damage, therefore one cannot know whether leukocyte plugged capillaries contributed to injury or perhaps were a result of it (and simply recruited). The idea that the diabetic endothelium is itself “sticky” and prone to clogging, perhaps as a result of chronic oxidative stress, is well appreciated in the diabetes literature ^56^ and could explain why most stalls were occluded with erythrocytes rather than leukocytes. Indeed there are reports in other diseases, such as thrombocythemia or the ApoE model of vascular dementia, where most stalled capillaries were composed of erythrocytes ^11, 57^. Therefore while leukocytes and in particular neutrophils, play a major role in stalling in mouse models of stroke and Alzheimer disease ^14, 18, 58^, their role may secondary to endothelial dysfunction in other conditions like diabetes.

Chronic inflammation is well appreciated as a contributing factor to capillary dysfunction in neurological disease and other organ systems. The present study as well as previous ones in rodents and humans, have shown that T1DM, especially if poorly controlled, is associated with elevated levels of inflammatory blood cytokines and chemokines ^59–62^. While we did not screen for all cytokines at both early and late time points, the fact that IL-10 levels were chronically high prompted us to investigate further. IL-10 is released by a variety of leukocytes and is elevated in retina and blood in both type 1 and 2 diabetes ^60, 63^. It is worth noting that IL-10 levels in are elevated in the blood of other conditions associated with systemic inflammation and perturbed microcirculation, such as LPS challenge, adverse reactions to CART therapy and even poor outcome from COVID-19 ^15, 16, 64^. One limitation with most previous studies was the correlational nature of IL-10 changes, thus whether it was serving a protective or pathogenic role was not well understood. To probe this issue further we employed orthogonal approaches to stimulate or inhibit IL-10 signalling through pharmacology and cell specific knockdowns. Contrary to our initial expectations that IL-10 would be protective as has been shown in other situations such as infection, stroke or endotoxin challenge ^27–30^, IL-10 signalling appeared to play a pathogenic role in capillary obstructions in diabetic mice. Consistent with this role, inhibiting IL-10 receptor signalling broadly with antibodies or genetic knockdown in endothelial cells significantly lowered the density of obstructed capillaries. The notion that IL-10R signalling could play a pathogenic role in certain disease states is not without precedent given that stimulating IL-10 in auto-immune diseases such as lupus or multiple sclerosis, can aggravate symptoms and sequelae ^65^. Studies have revealed that IL-10 deficiency inhibits Aβ clearance by microglia, worsening cognitive decline in mouse models of Alzheimer’s disease ^66^ ^67^. Indeed IL-10 signalling is complex and there are scattered reports showing it can promote inflammatory processes ^30^. For example, overexpression of IL-10 in the pancreas accelerated disease progression in non-obese diabetic mice by acting as a potent recruitment signal for leukocytes and damaging endothelial cells ^68^. Thus when viewed from the diverse IL-10 literature, our findings reinforce the notion that IL-10 signalling can play both protective and deleterious roles depending on the specific cell type, organ or condition/disease of study.

To our knowledge, this is the first study to show that blocking IL-10 signalling can exert a protective effect on the cerebral microcirculation and cognitive function, at least in the context of diabetes. Our qPCR data and cell specific knockdown experiments also indicate that activation of endothelial IL-10 receptors, which would be in direct contact with elevated IL-10 in diabetic blood serum, plays an important role in this phenomenon. The protective effects of inhibiting IL-10 signalling cannot be explained by a systemic change in blood glucose since levels were similar in all diabetic mice, regardless of treatment (see **Supp Fig. 3**). While IL-10 receptors are also expressed in other cell types like neutrophils and microglia, which have been implicated in regulating microvascular remodelling and blood flow in the diabetic retina ^69, 70^, these cells were not likely major contributors given that knockdown of IL-10 receptors specifically within neutrophils had no effect on capillary obstructions. Furthermore, the infection of cells following our viral therapy (using AAV.BR1.iCRE) was restricted to brain endothelium (see **Fig. 4c** showing cre-dependent reporter expression in vascular endothelium) and did not target microglia. However, since we did not manipulate IL-10 receptors in every possible cell type (only neutrophils and endothelium), we concede that IL-10 signalling in other cells, perhaps monocytes or lymphocytes, could be a factor in capillary stalling.

Our data raise the question of how treatments that target IL-10 signalling might work? Capillary stalling is a complex process that is regulated through multiple mechanisms including changes in blood flow or vascular tone, cell adhesion proteins, integrins in endothelial cells and leukocytes as well as alterations to the glycocalyx ^12, 71^. While testing each possibility is beyond the scope of the present study, our data show that treating diabetic mice with IL-10R blocking antibodies improve stimulus-evoked changes in cerebral blood flow and dilated lower order capillary branches off the penetrating arteriole by 25% (see **Figs. 5 and 6**). These findings suggest that endothelial IL-10 signalling could modulate vascular tone through astrocytes ^72, 73^ and/or contractile cells such as pericytes or smooth muscle cells ^74–76^. Indeed, a 25% increase in capillary diameter could greatly increase blood flow ^2^, which may be sufficient to push out erythrocyte plugged capillaries that manifest on the venous side. From a signalling perspective, there are reports showing that addition of IL-10 to cell culture can suppress vasoactive messengers such as nitric oxide, while IL-10 blocking antibodies can increase NO production ^77, 78^. IL-10 signalling could also augment the expression of cell adhesion or tight junction proteins which are up and down-regulated, respectively, in mice with elevated rates of capillary stalling ^12, 79^. For example, stimulating cultured endothelial cells with IL-10 enhances endothelial expression of VCAM-1, which is further potentiated in the presence of activated leukocytes ^80^. Permeability of the BBB could also be affected given that IL-10 can lower endothelial resistance and zona-occludins expression, while increasing permeability to sodium fluorescein ^81^. Thus, blocking IL-10 signalling could alleviate stalling through multiple avenues that future *in vivo* studies could explore and disentangle.

Previous studies have shown that alleviating capillary stalling with neutrophil or VEGF targeting antibodies rescues cognitive function in mice modelling Alzheimer disease ^12, 14^. In the present study we had to search for alternate therapeutic targets for diabetic mice because VEGF inhibition had no effect on capillary obstruction rates in our pilot experiments (unpublished data) and the majority of stalled capillaries in diabetic mice were not based on plugged leukocytes (∼25% based on leukocytes, see Fig. 1G). We therefore used the simplest and most clinically translatable approach of treating diabetic mice with an IL-10R neutralizing antibody for several weeks. This treatment significantly lower rates of capillary stalling (∼63% reduction) which also led to improved performance on behavioural tests of learning, memory and cognitive flexibility. These findings raise the question of how can blocking a small fraction of capillaries (∼1.82-3.64% in diabetic mice), appreciably affect cognition? Although our stalling estimates are largely comprised of short-lived stalls lasting at least 6.5s, experimental and computational data suggest that 2-4% of stalled capillaries can lead to a 5-20% decrease in cerebral blood flow, which also improved cognition in Alzheimer mice ^14, 82^. It is important to note that our microsphere experiments revealed that diabetes also increased the fraction of long-lived (∼3 day) capillary obstructions. Previous work from our lab has shown that long-lived capillary obstructions augment blood flow and diameter in connected capillaries for up to 2 weeks, and often resulted in vessel pruning which was not compensated for by sprouting of new capillaries ^10^. Further, capillary occlusion and subsequent vessel regression can lead to progressive degeneration of apical dendrites in cortical pyramidal neurons ^83^ or a reduction in local neural activity ^84^. Thus, it is reasonable to believe that greatly elevated numbers of short or long lasting capillary obstructions in diabetic mice could impact cognitive function and thereby represent a treatable target for ameliorating cognitive impairment.

## Methods

### Animals

Adult male and female mice (2-6 months of age) on C57BL/6J background were used in this study. For AAV mediated knockdown of IL-10 receptors in endothelial cells, we utilized *Il10ra* floxed mice (JAX# 028146) where loxP sites are expressed on exon 3 of the *Il10ra* gene ^85^. To manipulate IL-10 signalling in neutrophils, we crossed *Il10ra* floxed mice with the constitutive CRE driver mouse line *Mrp8/S100a8*-cre (JAX#021614). Previous work has shown *Mrp8/S100a8*-cre line has high specificity for neutrophils ^86, 87^. All mice were housed in groups on a 12h light/dark cycle in ventilated racks in a humidity (RH 40-55%) and temperature controlled room (21-23°C). Mice were provided food and water *ad libitum*. All experiments comply with the guidelines set by the Canadian Council on Animal Care and approved by the local university Animal Care Committee. Reporting of this work complies with ARRIVE guidelines.

### Induction of type 1 diabetes and insulin treatment

As previously described ^51^ type 1 diabetes was induced by two low dose injections of streptozotocin (i.p., STZ; Sigma# S0130; 75mg/kg per injection) dissolved in 50mM (pH 4.5) sodium citrate buffer over consecutive days. Mice that had received STZ injections but did not exceed the hyperglycemic threshold (>15mM) were given a single, additional dose of STZ. Non-diabetic (NDB) controls were administered sodium citrate buffer alone. Mice were given 5% sucrose water overnight after each day’s injection to prevent sudden hypoglycemia. Blood glucose levels were measured using glucometer (Accu-Chek, Aviva, Roche) every week by fasting mice for 2–3h and then withdrawing a drop of blood from the tail vein. To lower blood sugar levels in a subset of diabetic mice, one slow-release insulin pellet (0.1 U/24 h/implant, LinBit) was implanted subcutaneously under isoflurane anesthesia between the scapulae 1 week after confirmation of hyperglycemia. Blood glucose was tested weekly and additional insulin pellets were implanted if levels exceeded 15mM.

### Cranial window surgery

Mice were anesthetized with isoflurane (2% for induction and 1.3% for maintenance) in medical air (80% N_2_, 20% O_2_) at a flow rate of 0.7L/min. A temperature feedback regulator and rectal probe thermometer maintained body temperature at 37℃ throughout the procedure. After subcutaneous injection of lidocaine under the scalp, the scalp was cut across the midline and retracted. A custom metal ring for head fixing mice during imaging (∼1g in weight, outer diameter 11.3mm, inner diameter 7.0mm, height 1.5mm) was positioned over the right somatosensory cortex and secured to the skull with metabond adhesive. A circular area (diameter of ∼4-5mm) of skull within the metal ring was thinned using a high-speed dental drill, and ice-cold HEPES buffered artificial cerebrospinal fluid (ACSF) was periodically applied to the skull for cooling purposes. Fine forceps were used to remove the piece of skull and a 5 or 6mm circular coverslip was placed over the exposed brain and secured to the skull using cyanoacrylate adhesive. After the procedure, mice were injected with 0.03mL of 2% dexamethasone (i.p.) to reduce acute inflammation resulting from the procedure.

### Two-photon imaging and analysis of cortical microcirculation

*In vivo* two-photon imaging sessions began ∼6 weeks after cranial window surgeries and 4 weeks after diabetes induction (see **Fig. 1a**). Previous studies have shown that surgery induced inflammation and gliosis subsides by 3–4 weeks after window implantation ^88, 89^. Mice were lightly anesthetized with isoflurane (2% for induction, 1-1.5% for imaging) mixed in medical air. The metal ring on the mouse’s head was clamped into a metal bar in a custom-made frame to prevent movement related artifacts during imaging. Fluorescein or Texas Red labelled dextran (0.1mL of 1-5% of 70kDa dextran solution in saline; Sigma-Aldrich #46945 or Thermofisher D1830) was injected intravenously immediately before the start of imaging to visualize blood flow. For labelling and imaging of leukocytes ^36^, a subset of mice were intravenously injected with 0.1mL solution containing 0.4mg/kg FITC anti-mouse CD45.2 antibody (Clone 104, Biolegend, Cat #109806) and 1% Texas Red solution.

High-resolution two-photon image stacks of flowing and stalled capillaries were generated using an Olympus FV1000MPE multiphoton laser scanning microscope equipped with a mode-locked Ti:sapphire femtosecond laser (Mai Tai XF Deep See, Spectra-Physics). The laser was tuned to 800 or 850nm for excitation of Fluorescein or Texas Red dextran respectively, or 940nm for tandem imaging of FITC-CD45.2 labelled leukocytes and Texas Red dextran labelled blood plasma. Excitation power measured at the back aperture of the objective was typically between 17 and 60mW, and was adjusted to achieve similar levels of fluorescence within each imaging session. Images were acquired either with a 40× Olympus IR-LUMPlanFl water-immersion objective (NA = 0.8) or a 20× Olympus water-immersion objective (NA = 0.95), using Olympus FV10-ASW software. Three to four imaging areas per mouse were chosen based on proximity to the fore or hindlimb area of the somatosensory cortex. Images of the surface vasculature were used to re-locate the same imaging areas between sessions. During each imaging session, blood vessels were imaged to a depth of 200-300μm below the pial surface. Image stacks were collected at the following parameters depending on objective lens: a) 40× objective: 1.5μm Z-steps covering an area of 317.4×317.4μm (0.31μm per pixel) or b) 20× objective: 1.25μm Z-steps covering an area of 489.5×489.5μm (0.48μm per pixel). After imaging, mice were monitored while they recovered under supplementary heat, and then returned to their home cage ^90^.

Capillary stalls were identified by observers blinded to condition by first examining maximal Z-projection images (20-25 images) of cortical vessels. Capillaries that did not show any evidence of blood plasma streaks (streaks caused by cells moving through the lumen), as well as those that possessed dark gaps in the lumen (putative unlabelled RBCs) or FITC labelled leukocytes, were considered candidate stalls. These stalls were then followed up and verified by manually scrolling through 3-dimensional image stacks. A stalled capillary was determined if there was an absence of RBC streaks or presence of static cells (RBC or leukocyte) plugging the capillary for at least 3 imaging frames (minimum stall duration of 6.5s), similar to previously published criteria ^14^. To determine the branch order of stalled capillaries, vessels were traced back to the nearest penetrating arteriole or venule, with the aid of brightfield images of the cortical surface.

### Microsphere assay of capillary obstructions and analysis

As described in our previous studies ^5, 10^, we injected fluorescent microspheres into mice to examine experimental differences in susceptibility to capillary obstructions. Thus, male and female mice were briefly (<10 min) anesthetized with 1.5% isoflurane and intravenously injected with 3µL/g bodyweight of 5μm diameter fluorescent microspheres (1% solids; Bangs Laboratories, Inc., catalog# FCDG008). Importantly, experimental groups were run at the same time (eg. non-diabetic controls vs diabetic treatment groups) and injected from the same stock solution of microspheres to minimize cohort to cohort variability. Mice recovered under a heat lamp or heating pad immediately after injection before being returned to their home cage. Of note, we did not observe any abnormal behaviour or morbidity following injections. Three days after injection, mice were deeply anesthetized and the brain was extracted (without perfusion) and immersed in 4% PFA overnight before being transferred to 0.1M PBS. Brains were sectioned in the coronal plane on a Leica vibratome (T1000) at 50µm thickness. Every sixth section was mounted onto a gelatin-coated slide, and cover-slipped with Fluoromount G. Ultra-bright fluorescent microspheres were imaged on an upright wide-field Olympus BX51 microscope with a 2× Olympus Plan objective (NA = 0.05) using GFP excitation/emission filter sets on an Olympus DP73 digital CCD camera using CellSens software. Epi-fluorescent images were taken from approximately +2.7mm to −3.5mm from bregma, thereby covering much of the mouse forebrain. Using Image J FIJI (ver. 1.53q), regions of interest (ROI) were manually drawn over 13 different brain regions (see **Fig. 2**) we had previously conducted a detailed analysis of vascular parameters (length density, width, tortuosity; ^5^, guided by the Franklin and Paxinos Mouse Brain Atlas. Automated counting of microspheres across the mouse forebrain or within each region was achieved by thresholding pixels at of 67% maximum intensity. The density of capillary obstructions collapsed across the forebrain or within each brain region was expressed as # obstructions per cubic mm.

### Administration of IL-10 receptor neutralizing antibody and IL-10 protein

For inhibiting IL-10 receptor signaling *in vivo*, we randomly assigned diabetic C57BL/6J mice to receive an intravenous injection of either 250 µg/mouse of IL-10R blocking antibody (BioXcell #BE0050, *InvivoMAb* anti-mouse IL-10R, CD210, clone 1B1.3A) or equivalent volume of isotype control antibody (BioXcell #BE0088). Given the relatively durable activity of neutralizing antibodies, injections were always spaced out by 3-4 days (ie. 2 injections/week). To test whether IL-10 protein plays a role in promoting capillary obstructions, we intravenously injected non-diabetic mice with either recombinant mouse IL-10 protein (0.5µg per mouse; R&D #417-ML-005/CF) or albumin (0.5µg as control) over two consecutive days before injection of fluorescent microspheres, as described above.

### AAV based knockdown of endothelial IL-10Ra receptors

AAV-BR1-Cre vector was produced in Sf9 insect cells by the modified baculovirus expression system. Sf9 insect cells in Insect-XPRESS medium (Lonza, Basel, Switzerland) containing gentamycin (10 mg/liter) (Sigma-Aldrich, Darmstadt, Germany) was infected with high-titer stocks of two recombinant baculoviruses, one containing the AAV2-NRGTEWD (“AAV-BR1”) rep and cap genes ^41^ and the other containing a CAG promoter–driven iCre expression cassette embedded between AAV2 inverted terminal repeats based on the pAAV-CAG-iCre plasmid. Cells were cultured in normal atmosphere at 27°C and 110 rpm. AAV-BR1-iCre vector was purified by ultracentrifugation in a discontinuous iodixanol gradient, according to a previously published protocol. After ultracentrifugation, the purified AAVs contained in the 40% iodixanol fraction were aspirated and transferred to 10,000 molecular weight cutoff Vivaspin tubes (Sartorius, Göttingen, Germany) for dialysis in PBS. To generate mice with endothelial IL-10 receptor knockdown, we intravenously injected AAV-BR1-iCRE (20µl per mouse at 5.0×10^12^ GC/mL). The control group received comparable injections of AAV-BR1-eGFP. Three weeks after virus injection, mice were injected with fluorescent microspheres to assess susceptibility to capillary plugging as described above.

### Behavioural Testing

We performed a battery of behavioral tests starting at 4 and 8 weeks after induction of diabetes, with each testing period taking 1 week to complete (**Fig. 6a**). The 4 week time point was used to evaluate the initial impact of diabetes on behaviour and the 8 week time point was used to determine whether IL-10R blocking antibody treatment reversed any behavioural impairments.

#### Novel Object Recognition (NOR) Task

NOR was used to assess recognition memory ^45, 91^. The mice were first habituated to a plexiglass box (30 × 30 × 30 cm) devoid of any objects on habituation day for 5min. On the day of testing, mice were allowed to explore two identical objects, 12cm apart from each other, for 5min. The plexiglass box and the objects were wiped with 70%v/v ethanol to remove odors after each trial. Six hours later, a novel object was substituted for one of the familiar objects. The mice were again placed in the arena and left to explore the objects for 5 min. Using Ethovision software (version 11.5.1020, 2015; Noldus Information Technology), the amount of time spent exploring with its snout pointed in each object zone was determined as a percentage (time exploring novel object divided by total amount of time exploring objects). Similarly, preference ratio for the frequency of visits to novel zone were calculated.

#### Morris Water Maze

The Morris water maze was used to assess spatial learning and memory ^92^. The maze consisted of a water-filled circular pool (100cm diameter), and a hidden platform approximately 1cm below the surface. The pool was surrounded by curtains to shield the mouse from extraneous objects in the room and the experimenters during testing. Non-toxic white paint was added to the water to conceal the platform, and the temperature of the water was maintained at ambient room temperature. Mice were trained to locate the hidden platform by swimming using distinct visual cues placed around the maze. These cues varied in shape, and were kept constant throughout the experiment. EthoVision software (version 11.5.1020, 2015; Noldus Information Technology) was used to score escape latency (seconds) and swim speeds (cm/s) to the platform. The trial would begin with the mouse being placed in one quadrant facing the pool wall. The trial ended once the mouse located and stood on the platform for three seconds, or after a maximum of two minutes had elapsed without the mouse finding the platform. If the latter occurred, the experimenter placed the mouse on the platform, and allowed to it remain there for a minimum of five seconds. Each of the training sessions consisted of four trials, with each trial having a different starting point (southeast (SE), northwest (NW), northeast (NE), and southwest (SW)). Two training sessions were performed on each testing day, for a total of four training sessions over the two-day testing periods. Since the water maze was conducted twice in the same animals, the hidden platform was located in different quadrants for the 4 and 8 week testing sessions. Two days after mice completed the initial trials for learning the hidden platform location; a probe trial was performed to assess reference memory. This involved the removal of the hidden platform, and placement of the mouse into the maze for a two-minute period. The amount of time spent in each quadrant during this probe trial was recorded and a preference ratio was calculated by dividing the time spent in target quadrant by the time spent in other three quadrants. Three days after the probe trial test, we tested cognitive flexibility by switching the location of the hidden platform to a new location ^93^, which we refer to as the “platform reversal learning test”. Training sessions were run in the same manner as described above for initial platform learning.

#### Visible Platform Test

A visible platform test was performed in the Morris water maze at 8 weeks to determine if diabetes was associated with compromised vision. For this purpose, a platform that visually contrasted with the water was raised above the water line and the latency (s) to find the platform was recorded in the same manner as previously described.

#### Tape Removal Test

As previously described ^90, 94^, we assessed sensorimotor function of the forelimb using the adhesive tape removal test. For each session of testing, mice underwent three trials. Each trial was initiated by placing circular pieces of adhesive tape (5mm diameter) onto the palmar surface of both forepaws and placing the mouse into a clear glass cylinder. The mouse was filmed from below and allowed up to 2 min to remove the pieces of tape. The latency to remove each piece of tape was recorded using slow-motion video playback. Readings from left and right paw were averaged and presented in graphs as tape removal latency.

#### Open Field Test

To assess locomotive ability, mice were subjected to an open field test ^95^. The mouse was placed into the centre of a 100cm diameter circular arena, and the total distance travelled (cm) was recorded and tracked by Ethovision software. One trial of five minutes was conducted for each mouse at both 4 and 8 weeks.

#### Elevated Plus Maze

The elevated plus maze test was used to test anxiety-like behavior ^96^. This test involves a raised plus-shaped platform, with two arms with high borders (closed arms) and two that do not (open arms). Each mouse was run through one trial, which began when the mouse was placed in the centre of the elevated plus maze, and continued for five minutes. Ethovision software was used to track the mouse’s movements, and the percentage of time spent in the open arm was calculated. A higher open-arm percentage reflects lower anxiety in the mouse.

### Laser doppler measurements of cerebral blood flow

After behavioural testing was complete, laser doppler flowmetry was used to assess regional cerebral blood flow in response to stimuli known to increase blood flow ^97^. Mice were anesthetized with urethane (1.25g/kg) and the skull was carefully thinned over the right somatosensory cortex. The laser Doppler probe (Moor Instruments, MoorVMS-LDF1, PC version 2.2) was placed approximately 1mm above the skull with a stereotaxic arm. Three different stimuli were used to assess stimulus-evoked changes in blood flow: (i) inhalation of 5% CO_2_ mixed in medical air for 60s (x1 trial), (ii) vibrotactile hindlimb sensory stimulation (100Hz stimulation for 10s, x6 trials) (iii) 30s inhalation of 1% isoflurane (mixed in medical air; x1 trial). For each trial, we obtained a stable baseline before presenting each stimulus and recorded changes in blood flow. Using pCLAMP software, components of the cerebral blood flow response such as peak amplitude, latency to peak and area under the curve, were measured from 0-120s after CO2 exposure, 0-15s after sensory stimulation and 0-180s after isoflurane.

### Multiplex Immunoassay of immune cytokines and chemokines

Mice were deeply anesthetized with sodium pentobarbital and blood serum was collected. The blood was allowed to clot at room temperature for 10min and then spun at 1,500rpm for 10min in 4°C. The supernatant was collected, flash frozen in liquid nitrogen and stored at -80°C. On the day of testing, mouse serum samples were thawed on ice and were assayed using the Meso Scale Discovery (MSD) U-PLEX Biomarker Group 1 (mouse) 29-Plex panel assay (K15355K-1; lot# 393914-393917) as per manufacturer’s instructions to measure serum cytokine concentrations. The analysed cytokines and chemokines in the assay were IL-1β, IL-2, IL-5, IL-6, IL-10, IL-15, IL-17A, IL-17E/IL-25, IL-33, IP-10, KC/CXCL1, MCP-1, MIP-1α, MIP-2, MIP-3α, TNF-α. Samples were run in duplicates; plates were read using Quickplex SQ 120MM instrument and data were analyzed using the Discovery Workbench 4.0 software (Meso Scale Discovery). Briefly, the procedure includes several steps such as capture antibody coupling and plate coating; preparation of calibrator stocks and sample dilutions; addition of calibrators and samples, addition of detection antibody solutions; washing steps and reading the plates. Lower limits of calibrator detection (LLOD) for all assays were similar to product datasheets, demonstrating the sensitivity and robustness of the U-PLEX method. For serum sample signals within calibrator detection range, average calculated sample concentration % CV was 11%, showing the precision of the assays for quantifying changes in inflammatory response to diabetes.

### Endothelial cell separation

Mice were deeply anesthetized with 2% isoflurane and euthanized by decapitation. The cortex and cerebellum from the left hemisphere was immediately collected in 2mL RNase-free Microfuge Tubes (Thermo Fisher #AM12475) containing 1mL HBSS with no calcium and magnesium (Thermo Fisher #14175095). Brain tissues were cut into small pieces on a sterile 35mm petri dish using scalpel before transferring them into a 15mL falcon tube. The mechanical dissociation of brain tissue was performed using the Adult Brain Dissociation Kit (Miltenyi Biotec #130-107-677) as per manufacturer’s instructions. Briefly, cortex and cerebellum were homogenized by gently pipetting up and down ∼10 times with a 1mL pipette at 37⁰C. The recovered homogeneous cell mixture was gently applied to a smart strainer (Miltenyi Biotec #130-098-462) to remove the connective tissue. In the following steps, samples were always kept on ice unless otherwise indicated. Myelin was removed from the cell mixture using Myelin Removal Kit (Miltenyi Biotec #130-096-733) as per manufacturer’s instructions and then passed through LS columns (Miltenyi Biotec #130-042-401) using Quadro-MACS separator (Miltenyi Biotec #130-090-976). After 3 washes with Auto-MACS rinsing solution (Miltenyi Biotec #130-091-222) containing 0.5% BSA (Miltenyi Biotech #130-091-376), the flow through was centrifuged for 10min and re-suspended in Auto-MACS Rinsing Solution containing 0.5% BSA. Microglia were removed from the total cell suspension using CD45 microbeads (Miltenyi Biotec #130-052-301) by passing the cell mixture through LS columns. The flow through containing CD45-negative cells was then centrifuged for 10min. The cell pellet was re-suspended in Auto-MACS Rinsing Solution with 0.5% BSA and incubated with CD31 microbeads (Miltenyi Biotech #130-097-418) and passed through MS column (Miltenyi Biotech #130-042-201) using Octo-MACS separator (Miltenyi Biotec #130-042-108). After 3 washes with Auto-MACS rinsing solution with 0.5% BSA, column bound CD31 positive endothelial cells were collected by adding 1mL Auto-MACS rinsing solution with 0.5% BSA. The collected sample volume was centrifuged at max speed in a benchtop centrifuge for 10 minutes and after discarding the supernatant, the endothelial cell pellet was flash frozen and kept at -80⁰C until further use.

### RNA extraction and quantitative real-time polymerase chain reaction

Total RNA was extracted using RNeasy Mini kit (QIAGEN #74104) with deoxyribonuclease (Qiagen #79254) treatment step following manufacturer’s instructions either from flash frozen brain tissue or isolated cells. RNA purity and concentrations were estimated using a NanoDrop 1000 Spectrophotometer (Thermo Fisher Scientific, Waltham, MA, USA) and samples were stored at - 80⁰C until further use. cDNA was prepared using high-capacity cDNA synthesis kit (Applied Biosystems #4368814) from fifty nanograms RNA with the help of Bio-Rad T100 thermocycler (Bio-Rad, Mississauga, Ontario, Canada). cDNA was diluted 5-fold for reverse transcription quantitative polymerase chain reaction (RT-qPCR) analysis. Primer specificities were calculated both online using NCBI primer blast and experimentally using melt curve analysis. A 5-point 10 fold serial dilution was used to test primers efficiency and those that achieved efficiencies between 90 - 110% and were used for gene expression analysis. RT-qPCR was carried out using 10μL reaction mixture containing 1μL cDNA, 0.5μL of each primer (10μm stock), 3μL RNase, DNase free water and 5μL of SYBR Green Master Mix (Applied Biosystems). Thermocycling conditions used were 50⁰C (2 min), 95⁰C (2 min), followed by 40 cycles of 95⁰C denaturation (15 sec), 60⁰C /62⁰C annealing (1 min). Fluorescent signals were acquired by StepOne plus system and data were analysed by Design and Analysis Software Version 2.4.3 (Applied Biosystems). Triplicate reactions were performed for each sample and Ct values were averaged and normalized to the geometric mean of TBP and HPRT using comparative delta-delta Ct (ΔΔCt) method to calculate relative mRNA levels. Forward and reverse primers used for RT-qPCR were as follows: *Tbp* (housekeeping): CCCCACAACTCTTCCATTCT and GCAGGAGTGATAGGGGTCAT; *Hprt* (housekeeping): AGCCTAAGATGAGCGCAAGT and TTACTAGGCAGATGGCCACA; *Il10ra (exon 3 specific)*: AACCTGGAATGACATCCATATC and CCACTGTGAAGCGAGTCTCAGT; *Vegfr2*: TTTGGCAAATACAACCCTTCAGA and GCAGAAGATACTGTCACCACC; *Tie2*: GAGTCAGCTTGCTCCTTTATGG and AGACACAAGAGGTAGGGAATTGA; *Cd31*: ACGCTGGTGCTCTATGCAAG and TCAGTTGCTGCCCATTCATCA; *Cdh5*: CCACTGCTTTGGGAGCCTT and GGCAGGTAGCATGTTGGGG; *Tmem119*: CTTCACCCAGAGCTGGTTCCATA and CCGGGAGTGACACAGAGTAG; *Cd45*: GTTTTCGCTACATGACTGCACA and AGGTTGTCCAACTGACATCTTTC; *NeuN*: ATCGTAGAGGGACGGAAAATTGA and GTTCCCAGGCTTCTTATTGGTC.

### Statistics

All data were statistically analysed using GraphPad Prism (versions 7 and 8, RRID:SCR_002798). Student’s *t* tests (paired or unpaired, as appropriate) were used to analyze simple between group differences (ie. when only 2 or 3 groups compared on a single factor). Two-way ANOVAs were used to compare differences between treatment groups across time, testing session or brain region. Significant main effects from ANOVAs were analyzed with Tukey’s multiple comparisons tests. *P*<0.05 was considered statistically significant. All experimental values are presented as means ± SEM.

### Data Availability

Data generated for this study are available from the corresponding author upon reasonable request.

## Acknowledgements

We are grateful to Pat Reeson and Kerry Delaney for their input, as well as Angie Hentze and Taimei Yang for managing the mouse colony. We also thank David Attwell (UCL) for thoughtful discussions regarding the work. This research was supported by operating, salary and equipment grants to C.E.B. from the Canadian Institutes of Health Research (CIHR), Heart and Stroke Foundation (HSF), Natural Sciences ands Engineering Research Council (NSERC). S.S. was supported by a post-doctoral fellowship from Michael Smith Foundation for Health Research.

## Author Contributions

C.E.B, and S.S. conceived the study and wrote the manuscript. S.S, K.A.T., M.C., R.B., A.P.C., T.P.B, R.D.F, L.A.R., J.K., and C.E.B. provided key intellectual insights, reagents, performed experiments and/or analyzed data.

