## Supplementary data for "A pathogenic role for IL-10 signalling in capillary stalling and cognitive impairment in type 1 diabetes"

**Supplementary Figure 1**

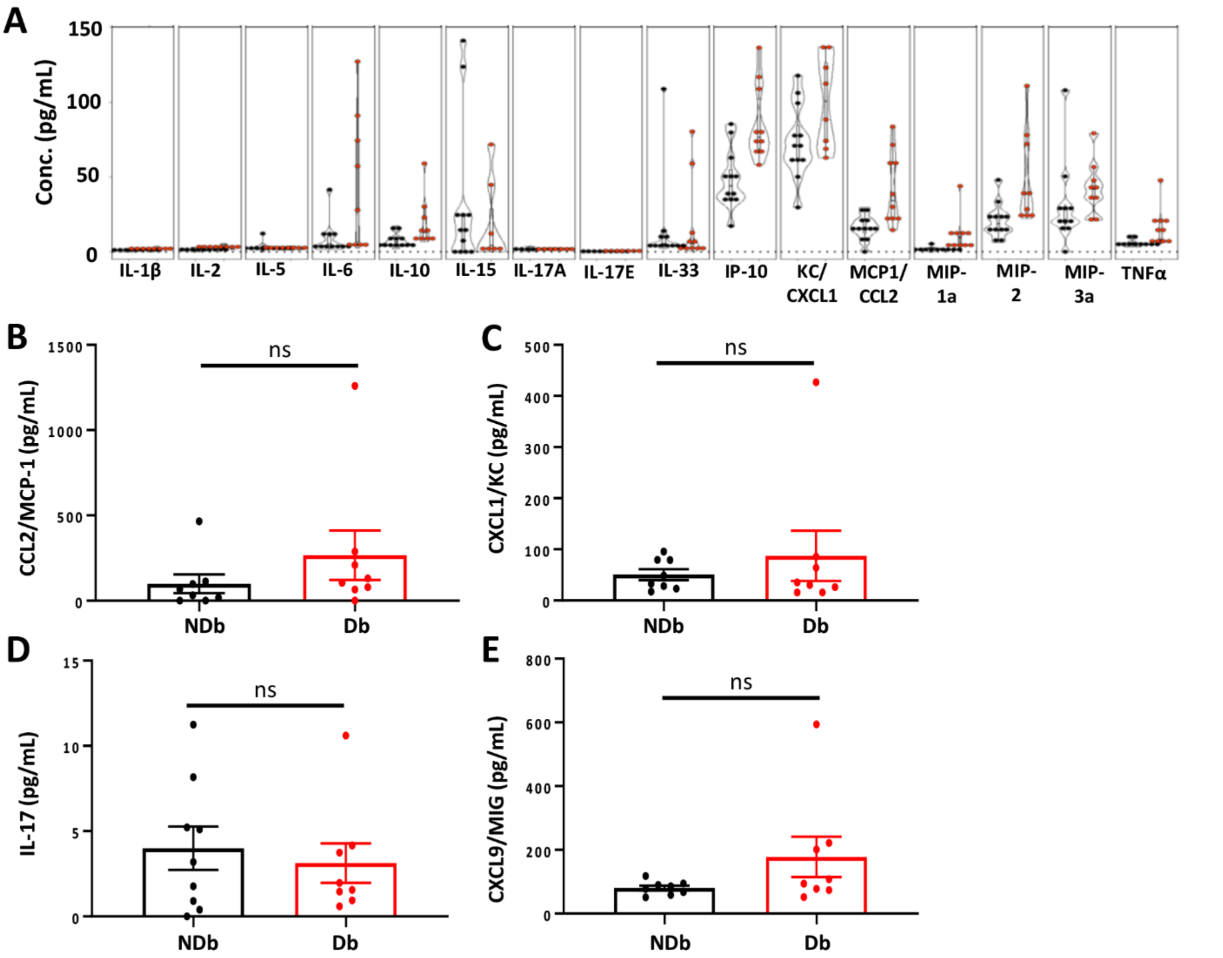


**Supplementary Figure 1. Blood serum cytokine levels in non-diabetic and diabetic mice.** (**A**) Concentration for each cytokine/chemokine (pg/mL) in blood serum of non-diabetic (black dots) and diabetic mice (red dots) at 8 weeks. (**B-E**) Graphs show expression of CCL2/MCP-1 (**B**), CXCL1/KC (**C**), IL-17A (**D**), and CXCL9/MIG (**E**) in blood serum 4 weeks after confirmation of hyperglycemia. Data analysed by unpaired two-tailed t-test. Error bars: mean ± SEM.

### Supplementary Figure 2

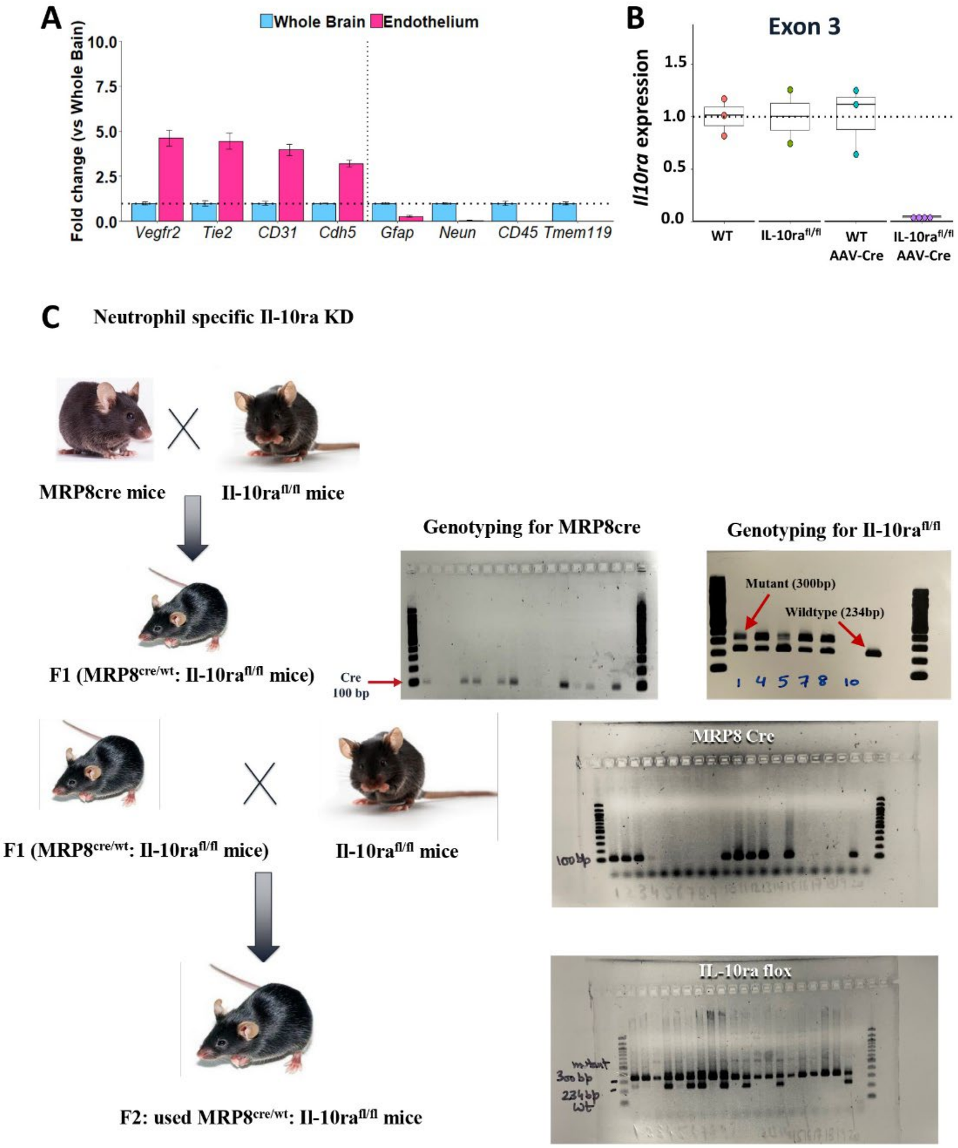


**Supplementary Figure 2. Validation of *IL10ra* knockdown in endothelial cells and neutrophils.** (**A**) Isolated endothelial cells showing enriched expression of genes typically expressed in endothelium (*Vegfr2, Tie2, Cd31, Cdh5*), with very low levels of gene expression associated with astrocytes (*Gfap*), neurons (*NeuN*), leukocytes (*Cd45*) or microglia (*Tmem119*). (**B**) qPCR data from endothelial cells show loss of Exon 3 *IL10ra* gene expression in endothelial cells isolated from *Il10ra floxed* mice injected with AAV-BR1-iCRE (n=4 mice), relative to controls (WT: Wild type; *Il10ra flox/flox*: *Il10ra flox/flox* mice; WT mice injected with AAV-BR1-iCRE). (**C**) Breeding strategy and PCR results for *Mrp8cre:IL10ra* floxed mice.

**Supplementary Figure 3**

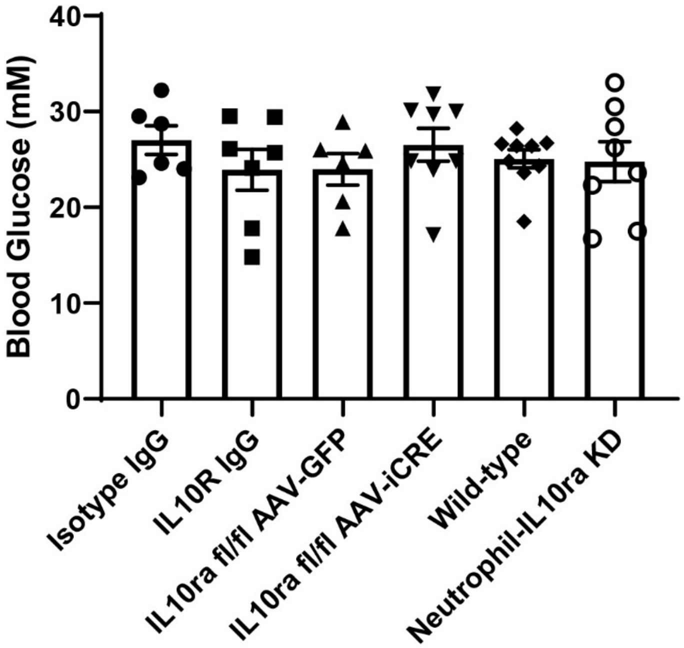


**Supplementary Figure 3. Blood glucose levels in diabetic mice at 4 weeks (for experiments shown in Fig 4) across different treatment groups.** One-way ANOVA indicated no significant differences between diabetic groups**.**

**Supplementary Figure 4**

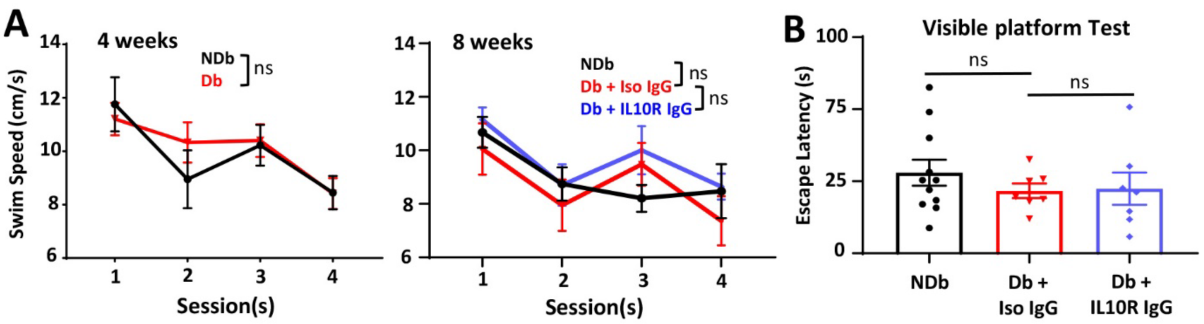


**Supplementary Figure 4. Effect of diabetes on swim speed and visibility of mice.** (**A**) Swim Speed in Morris water maze at 4 and 8 week testing periods (n=11, 15, 12, 7 and 8 mice, respectively). (**B**) Escape latencies in the visible platform test at 8 week testing period (n=11, 7 and 8 mice, respectively). Data analysed with two-way ANOVA (A) and unpaired two-tailed t-tests (B). Error bars: mean ± SEM.
